# Fibroblasts carrying intermediate *C9orf72* hexanucleotide repeat expansions from iNPH patients show impaired energy metabolism but no cell pathologies

**DOI:** 10.1101/2024.05.28.595117

**Authors:** Dorit Hoffmann, Ville Korhonen, Hannah Rostalski, Nadine Huber, Sami Heikkinen, Tomi Hietanen, Rebekka Wittrahm, Stina Leskelä, Päivi Hartikainen, Tuomas Rauramaa, Eino Solje, Anne M. Portaankorva, Mikko Hiltunen, Ville Leinonen, Annakaisa Haapasalo

## Abstract

Long *C9orf72* hexanucleotide repeat expansions (C9-HRE) are the most common genetic cause of frontotemporal dementia (FTD), a group of neurodegenerative syndromes leading to cognitive dysfunction and frontal and temporal atrophy. FTD is a potential comorbidity of idiopathic normal pressure hydrocephalus (iNPH) and carrying the C9-HRE can modify the age-of-onset in iNPH patients. While intermediate-length C9-HRE (<30 repeats) are often considered non-pathogenic, the exact pathological cutoff is unclear. In this study, we assessed whether fibroblasts from iNPH patients carrying intermediate C9-HRE display C9-HRE-associated pathological hallmarks and changes in cellular function. C9-HRE-associated RNA foci were not detected in the intermediate carriers. The number of p62-positive puncta was significantly increased only in long C9-HRE carrier fibroblasts, in line with p62-positive intracellular inclusions observed in a brain biopsy from the patient. Specific parameters of mitochondrial respiration were significantly reduced in both the long and intermediate C9-HRE carrier fibroblasts. Fibroblasts from the intermediate C9-HRE carriers showed upregulated glycolytic activity, possibly to counteract the reduced mitochondrial respiration, which could not be observed in the long C9-HRE carriers. In conclusion, these data suggest that while the long C9-HRE leads to more severe cellular pathologies than intermediate C9-HRE, the latter might predispose cells to pathological changes.

## 1 Background

Frontotemporal dementia (FTD) is a progressive early-onset (<65 years) neurodegenerative disorder, characterized by degeneration in the frontal and temporal lobes of the brain. Behavioral variant frontotemporal dementia (bvFTD) is the most common clinical subtype of FTD. Approximately half of the FTD cases are caused by mutations in different genes, including *MAPT* (microtubule associated protein tau) and *GRN* (progranulin), or by long GGGGCC hexanucleotide repeat expansions in *C9orf72* (C9-HRE), the most common genetic cause underlying both familial and sporadic FTD and amyotrophic lateral sclerosis (ALS) [1–7]. In the affected individuals, the C9-HRE length can reach several hundreds or thousands of copies.

Idiopathic normal pressure hydrocephalus (iNPH) is a neurological disease characterized by a group of three clinical findings (Hakim’s triad), which are gait disturbances, cognitive impairment, and urinary incontinence [8]. The patients might show all or some features of the triad and can additionally display impaired frontal executive function [9] and enlarged ventricles in clinical imaging [10,11]. Similar to bvFTD, iNPH can present symptoms such as decline in executive function, psychomotor slowness, and behavioral and personality changes [12]. In fact, even though the most common comorbidities of iNPH are hypertension, Alzheimer’s disease (AD), and vascular dementia [10,13], FTD has also been described as a comorbidity for iNPH [10,14]. Interestingly, long C9-HRE can modify the age-of-onset in iNPH patients as shown in a cohort of Finnish iNPH patients [15].

While C9-HRE with intermediate repeat lengths of less than 30 typically do not cause FTD and are often considered non-pathogenic [16], there is some evidence that they could be associated with ALS and other neurodegenerative diseases. Two meta-analyses have found an association between intermediate repeats of 24-30 and ALS, and the authors suggest that repeats of 24 or longer should be considered pathogenic [17,18]. For shorter expansions, the results are contradictory. A study in Finnish patients found that carrying intermediate-length alleles increases the risk of ALS when one of the alleles has ≥ 17 repeats [19], but this could not be observed in a larger study [20]. A significant association between clinically diagnosed Parkinson’s disease (PD) and the intermediate C9-HRE has been reported [21]. This association, however, could not be corroborated in a cohort of autopsy-confirmed PD cases [22]. Moreover, a possible association between the intermediate C9-HRE and atypical Parkinsonian syndromes has been suggested in a few studies containing a small number of patients [23,24]. Cali *et al*. [25] assessed whether the intermediate C9-HRE could be a genetic risk factor for corticobasal degeneration (CBD), a neurodegenerative disease in the FTD spectrum that shares similarities with PD but displays tau protein brain pathology. They found that the number of individuals with the intermediate C9-HRE was significantly higher among CBD cases as compared to controls.

The main pathological mechanisms associated with the long C9-HRE are gain-of-toxic function through accumulation of RNA foci and production of dipeptide repeat (DPR) proteins (poly-GP, poly-GA, poly-GR, poly-PA, and poly-PR) and loss-of-function due to haploinsufficiency leading to reduced C9orf72 mRNA and protein levels [26–29]. While these pathological hallmarks are specific for the long C9-HRE, additional hallmarks have also been detected in the CNS of FTD and ALS patients with and without the C9-HRE, such as intracellular inclusions formed by the accumulated TAR DNA-binding protein-43 (TDP-43) or sequestosome 1 (p62/SQSTM1, hereafter p62) [30–34]. Data on pathological changes in the intermediate C9-HRE carriers are sparse. It has been reported that the intermediate C9-HRE does not lead to the formation of RNA foci or DPR proteins. Furthermore, increased C9orf72 mRNA and protein levels were detected in patients and CRISPR/cas9 knock-in iPSC-derived neural progenitor cells carrying intermediate C9-HRE [25]. In the same study, the authors showed that C9orf72 overexpression in HeLa cells expressing a single copy of long C9-HRE affected autophagic function by promoting autophagy under nutrient-rich conditions while impairing autophagy during starvation-induced stress. Current data also suggest that both C9-HRE-associated gain-of-toxic-function and loss-of-function can impair mitochondrial function. Defective autophagy and production of DPR proteins may affect the energy metabolism of cells. Mitophagy, a special form of autophagy, is responsible for the elimination of damaged mitochondria [35] and the failure in this mitochondrial quality control mechanism can lead to mitochondrial dysfunction, which has been described in both FTD and ALS [36]. Moreover, impaired mitochondrial function has been reported in iPSC-derived motor neurons of patients carrying the C9-HRE [37].

While research on FTD and ALS-related pathologies has naturally focused mostly on neurons in FTD and ALS patient brain and iPSC-derived neurons, there are some studies showing that peripheral cells can also display distinct pathologies related to the C9-HRE and in general to ALS and FTD. RNA foci and poly-GP and poly-GA proteins have been detected in skeletal muscle biopsies from ALS patients carrying the C9-HRE [38]. iPSC-derived myocytes from C9-HRE-carrying ALS patients displayed RNA foci and expressed the poly-GR protein [39,40]. RNA foci have also been previously described in C9-HRE-carrying ALS or FTD patient-derived skin fibroblasts by us and others [41,42]. Other cellular pathologies have also been previously observed in fibroblasts. For example, increased levels of p62 and LC3II have been detected in C9-HRE-carrying ALS/FTD patient-derived fibroblasts, suggesting defective autophagy under stress conditions [43]. In our previous study, we observed p62 accumulation but no changes in basal or induced autophagy in both C9-HRE-carrying and non-carrying FTD patient fibroblasts [42]. Moreover, fibroblasts from patients with sporadic ALS and ALS patients carrying mutations in *VCP*, *SOD1*, or *TARDBP* genes have been reported to display impaired mitochondrial function [44–46]. Work by us and others in fibroblasts from ALS and FTD patients carrying the C9-HRE also showed mitochondrial dysfunction [47]. Taken together, these studies show that cells other than neurons, such as skin fibroblasts, can display some of the pathological hallmarks and altered cellular functions connected to the C9-HRE. These patient-derived cells might therefore be suitable for testing the effects of therapeutic interventions targeting specific pathways or in biomarker research in the future.

Based on these current data, we aimed to assess whether the intermediate C9-HRE have effects on cellular pathologies and function similar to those previously observed in the long C9-HRE carriers. To this end, we focused on characterizing skin fibroblasts from iNPH patients carrying intermediate C9-HRE. This is of particular interest as the pathogenic cutoff of the intermediate repeat length has been under debate and the cellular effects of the intermediate repeats have not been studied in much detail previously.

## 2 Material and Methods

### 2.1 Study subjects, skin biopsies, ethical permits, and genotyping

Skin punch biopsies were obtained at Neuro Center, Neurology, Kuopio University Hospital, Kuopio, Finland. Five iNPH patients, of whom four were intermediate carriers (10 to 23 repeats) of the C9-HRE and one long C9-HRE carrier (>60 repeats), as well as two FTD patients carrying the long C9-HRE, and three age-matched healthy controls were included in the study cohort. Both males and females were included in the cohort. Only one long C9-HRE-carrying iNPH patient could be included because these patients are rare. The C9-HRE carriership status in these individuals was confirmed from blood samples by repeat-primed PCR [20]. Brain biopsy samples from the iNPH patients had been previously assessed and p62-positive inclusions were found in one of the intermediate C9-HRE carriers and the long C9-HRE carrier. All the participants gave a written informed consent. The study was performed according to the Declaration of Helsinki. The research protocol has been approved by the Research Ethics Committee of the Northern Savo Hospital District, (currently: Medical Research Ethics Committee of Wellbeing Services County of North Savo) Kuopio, Finland (ethical permits 16/2013, 254/2015 and 276/2016). Skin biopsy samples were pseudonymized and handled using code numbers.

### 2.2 Culturing and treatments of fibroblasts

Fibroblasts were obtained from skin biopsy samples as described previously [42]. The fibroblasts were cultured in Iscove’s Modified Dulbecco’s Medium (IMDM, 21980032, Gibco) with 20% heat inactivated fetal bovine serum (FBS, 10270106, Gibco), 1x MEM Non-Essential Amino Acids (11140050, Thermo Fisher) and 100 U/ml penicillin and 100 µg/ml streptomycin (15140122, Thermo Fisher) (= fibroblast medium) at +37°C and 5% CO2.

For autophagy induction, cells were treated with 200 nM of Torin 1 (4247, Tocris) for 24h. To assess basal autophagy, cells were treated with 300 nM bafilomycin A1 (BafA1, B1793, Sigma-Aldrich) for 6 h to block the late phase of autophagy. To block protein degradation through the ubiquitin-proteasome system (UPS), 10 µM lactacystin (Enzo Life Sciences) was used for 16h [48]. Dimethyl sulfoxide (DMSO, D2650, Sigma-Aldrich) was used as a vehicle control.

### 2.3 Immunocytochemistry

For immunocytochemistry, glass coverslips were coated with 0.3% gelatine for 30 min at +37°C in 24 well plates. Fibroblasts were plated at a density of 20,000 cells/well in a 24 well plate and fixed after 24 h in 4% paraformaldehyde (PFA, 28908, Thermo Scientific) for 10 min at room temperature (RT). Cells were permeabilized with 0.1% Triton X-100 (X100, Sigma-Aldrich) for 10 min at RT and blocked for 30 min at RT in 1% bovine serum albumin (BSA, A9647, Sigma-Aldrich). For overnight incubation at +4°C the following primary antibodies were used: anti-TDP-43 (1:100, 10782-2-AP, Proteintech), anti-phospho-TDP-43 (1:200, CAC-TIP-PTD-M01, CosmoBio) and anti-p62 (1:200; sc-28359, Santa Cruz). The coverslips were incubated for 1 h at RT with one of the following secondary antibodies: goat anti-rabbit Alexa Fluor® 488 (1:500, A-11008, Invitrogen) was used for TDP-43, goat anti-mouse Alexa Fluor® 488 (1:500, A-11029, Invitrogen) for phospho-TDP-43 and goat anti-mouse Alexa Fluor® 568 (1:500 A11004, Invitrogen) for p62. Coverslips were mounted with Vectashield Vibrance antifade mounting medium containing 4′,6-diamidino-2-phenylindole (DAPI) (H-1800, Vector Laboratories) for immunocytochemistry with p62 or, for immunocytochemistry with TDP-43 and phospho-TDP-43, with a 1:1 mix of mounting medium with DAPI and Vectashield Vibrance antifade mounting medium with TRITC-Phalloidin (H-1600, Vector Laboratories). Images were taken with an Olympus BX51 microscope and analyzed with ImageJ (version 1.52 p, Fiji, NIH).

### 2.4 Immunohistochemistry

From the iNPH patients, right frontal cortical brain biopsy was obtained during insertion of the ventricular catheter for CSF shunt. Cortical biopsies were collected using biopsy forceps or a needle prior to the insertion of an intraventricular catheter for 24-hour monitoring of intracranial pressure or shunting. The biopsies were fixed in buffered formalin and embedded in paraffin. The resulting 7-μm sections were processed using standard techniques, including deparaffinization and rehydration. All sections were then stained using haematoxylin-eosin (H&E) and immunohistochemical methods. After pretreatment, the sections were blocked using normal goat serum for 30 minutes to reduce non-specific reactions. Epitopes were then unmasked and p62 antibody (1:1000, 610832, BD Biosciences) was added to the sections, which were incubated overnight at 4°C. The next day, the sections were incubated with a biotinylated secondary antibody and then with a streptavidin enzyme conjugate (85-8943, LABSA Zymed Laboratories) at room temperature for 30 minutes to visualize the reaction products. Immunostained sections were counterstained with Harris’ haematoxylin, dehydrated, and mounted in DePex (BDH Chemicals, Hull, UK). An experienced neuropathologist evaluated all sections using light microscopy. p62 immunohistochemistry was classified as present or absent.

### 2.5 Fluorescence *in situ* hybridization (FISH)

FISH was performed using a protocol based on a previous publication [29], with some modifications. Cells on gelatine-coated coverslips were fixed with 4% PFA in diethyl pyrocarbonate (DEPC)-PBS, permeabilized with 0.2% Triton X-100/DEPC-PBS, washed twice with DEPC-PBS and then incubated twice for two minutes in 70% ethanol and once in 100% ethanol for 2 minutes. This was followed by incubation in hybridization buffer (10% dextran sulfate, 50% formamide, 50 mM sodium phosphate buffer (pH 7), 2 x SSC) at 55°C for 30 min. Prior to use, the locked nucleic acid (LNA) probe TYE^TM^ 563-(CCCCGG)_3_ (Exiqon; recognizing the expanded G_4_C_2_ repeats) and the TYE^TM^ 563-(CAG)_6_ negative control probe (Exiqon) were denatured at 80°C for 5 min and diluted to 40 nM with hybridization buffer. The hybridization of the samples with either probe was performed in a light-protected chamber at +55°C for 3 h. Confocal images were acquired with LSM800 (Zeiss) microscope.

### 2.6 p62 puncta analysis

Immunocytochemistry was performed with the p62 antibody and DAPI as described above. For p62 puncta analysis, the number of cells per image was calculated using DAPI images and p62 images were converted into binary images and puncta of a defined size were used for further analysis. Images of cells stained without primary antibody were used for background subtraction and thresholding. The mean size of p62 positive puncta was calculated per image and mean number of p62 positive puncta per cell was calculated by dividing the number of puncta per image with the number of cells per image. The intensity was quantified as sum intensity and then normalized to the puncta size as described previously [42].

### 2.7 Protein extraction from cells and Western blotting

Proteins were extracted in lysis buffer (10 mM Tris-HCl, 2 mM EDTA, 1% SDS) containing protease and phosphatase inhibitor (1862209 and 1862495, Thermo Scientific). To measure protein concentration, bicinchoninic acid assay (BCA, 23225, Thermo Scientific) and a plate reader (Infinite® M200, Tecan Group Ltd.) were used. Eight µg of protein were loaded on sodium dodecyl sulfate–polyacrylamide gel electrophoresis (SDS-PAGE) gels (NuPAGE Novex 4-12% Bis-Tris mini or midi, NP0335 or WG1402BOX, Invitrogen) and run for 1 h 40 min at 100 V. With a Trans-Blot® TurboTM Transfer System (Bio-Rad, 25 V, 1.0 A, 30 min), proteins were transferred on 0.2 µm polyvinylidene fluoride (PVDF) membranes (1704157, Bio-Rad). Unspecific binding was blocked with 5% non-fat dry milk or bovine serum albumin (BSA A9647, Sigma-Aldrich) in 1 x Tris-buffered saline with 0.1% Tween 20 (93773, Sigma-Aldrich) (TBST) for 1 h at RT. The protein bands were detected by incubating the membrane with primary antibodies (see below) overnight at +4°C and horse radish peroxidase-conjugated secondary antibodies (1:5000, NA934 or NA931, GE Healthcare) for 1 h at RT. Proteins were detected with enhanced chemiluminescence (ECL) detection reagents (RPN2236 or RPN2235, Amersham Biosciences, GE Healthcare,) and ChemiDoc™ XRS+ System (Bio-Rad). Intensities of the detected protein bands were quantified with Image Lab™ software (6.0.1, Bio-Rad). Membranes were stripped with stripping buffer (21063, Thermo Scientific) for 10 min at RT, washed in 1 x TBST and re-probed with other antibodies. The following primary antibodies were used: anti-pULK1Ser757 (1:1000, # 14202S, Cell Signaling Technology), anti-ULK1 (1:1000, #8054, Cell Signaling Technology), anti-C9orf72 (1:500, 22637-1-AP, Proteintech), anti-SQSTM1/p62 (#5114, 1:1000, Cell Signaling Technology), anti-LC3B (1:3000, ab51520, Abcam), anti-poly-ubiquitinated proteins (FK1, 1:1000, BML-PW8805-0500, Enzo Life Sciences), anti-TDP-43 (1:1000, 10782-2-AP, Proteintech), anti-phospho-TDP-43 (1:1000, TIP-PTD-P02, CosmoBio) and anti-beta-actin (1:1000, ab8226, Abcam). The data are shown as median ± interquartile range or mean ± standard error of the mean (SEM). The protein levels were normalized to the levels of β-actin in the same sample and this ratio was set to 100 in (vehicle-treated) control cells. The protein levels are shown as % compared to those in vehicle-treated control cells (set to 100%).

### 2.8 Energy metabolism (mitochondrial respiration and glycolysis)

For the experiments on the energy metabolism, fibroblasts were plated (5000 cells/well) in uncoated Seahorse XF96 Cell Culture Microplates (101085-004, Agilent) with 8 wells per cell line in each experiment. For normalization of the data, cells were stained with Vybrant™ DyeCycle™ Green Stain (5 µM, V35004, Thermo Fisher) after completing the Cell Mito Stress Test or Glycolysis Stress Test. Images were acquired with 4x objective from brightfield and the IncuCyte® S3 (Essen BioScience). IncuCyte® software (v2019B) was used to count the number of cells per well in the green fluorescence channel. Parameters were calculated using the Wave 2.6.0 software (Agilent) and results were normalized to the number of cells counted per well

#### 2.8.1 Mito Stress Test

Mito Stress Test was performed 48 h after plating using assay parameters provided by Agilent. On the day of the experiment, medium was changed to Seahorse XF DMEM medium (103575-100, Agilent) supplemented with 10 mM Seahorse XF glucose solution, 2 mM Seahorse XF L-glutamine solution and 1 mM Seahorse XF pyruvate solution (103577-100, 103579-100 and 103578-100, all from Agilent) and cells were kept in a CO_2_ free incubator for 45 min prior to starting the Cell Mito Stress Test. The following final concentrations of electron transport chain modulators were used: 2 µM carbonyl cyanide-4-(trifluoromethoxy)phenylhydrazone (FCCP), 1 µM oligomycin and a mixture of 1 µM antimycin A and 1 µM rotenone (C2920, 75351, A8674 and R8875, all from Sigma-Aldrich). With a Seahorse XFe96 analyzer (Agilent), changes in oxygen consumption rate (OCR) in response to injections were detected. First, basal respiration is measured and then oligomycin, which blocks complex V (ATP synthase), is added. The subsequent decrease in OCR is linked to cellular ATP production. Adding the uncoupling agent FCCP leads to a collapse of the proton gradient, causing uninhibited electron flow through the ETC and oxygen consumption by complex IV reaches the maximum. With the OCR following FCCP injection (maximal respiration), also the spare capacity can be calculated, which indicates the celĺs ability to respond to an increased energy demand. The mitochondrial respiration is shut down completely with the injection of rotenone and antimycin A (they block complexes I and III, respectively), allowing the calculation of nonmitochondrial respiration driven by processes outside the mitochondria [49].

#### 2.8.2 Glycolysis Stress Test

Glycolysis Stress Test was performed 48 h after plating using assay parameters provided by Agilent. On the day of the experiment, medium was changed to Seahorse XF DMEM medium (103575-100, Agilent) supplemented with 2 mM Seahorse XF L-glutamine solution (103579-100, Agilent) and cells were kept in a CO_2_ free incubator for 1 h prior to starting the Glycolysis Stress Test. For the experiments, the following final concentrations were used: glucose 10 mM (103577-100, Agilent), oligomycin 1 µM (75351, Sigma-Aldrich) and 50 mM 2-Deoxy-D-glucose (2-DG, D6134-5G, Sigma-Aldrich). Changes in extracellular acidification rate (ECAR) in response to injections were detected with Seahorse XFe96 analyzer (Agilent). Prior to the assay, the cells were kept in glucose-free medium. Before the first injection, the non-glycolytic acidification is measured, since at this point the cells do not perform glycolysis because the medium does not contain glucose. The first injection adds a saturating amount of glucose and by measuring the ECAR, the rate of glycolysis under basal conditions is assessed. The next injection, oligomycin, inhibits the mitochondrial ATP production, so energy production is shifted to glycolysis. The increase in ECAR following this shift shows the maximal glycolytic capacity from which the glycolytic reserve can also be calculated. The final injection of 2-DG inhibits glycolysis by competitive binding to glucose hexokinase. The decrease in ECAR following this injection, confirms that the changes in ECAR observed during the experiment are due to glycolysis

### 2.9 RNA extraction and global RNA sequencing

Total RNA was isolated (11828665001, Roche Molecular Systems, Inc.) according to the manufacturer’s instructions and RNA concentrations were measured using NanoDropTM One (Thermo Scientific).

Bulk RNA sequencing (RNA-seq) was performed using RNA extracted as described above. Library preparation and RNA sequencing was conducted by Novogene (UK) Company Limited. In brief, mRNA enrichment was performed with oligo(dT) bead pulldown, from where pulldown material was subjected to fragmentation, followed by reverse transcription, second strand synthesis, A-tailing, and sequencing adaptor ligation. The final amplified and size-selected library comprised of 250-300 bp insert cDNA. The paired-end, 150 bp sequencing was executed with an Illumina high-throughput sequencing platform. Sequencing yielded 20.8–28.1 million sequenced fragments per sample.

The 150 nt pair-end RNA-seq reads were quality controlled using FastQC (version 0.11.7) (https://www.bioinformatics.babraham.ac.uk/projects/fastqc/). Reads were trimmed with Trimmomatic (version 0.39) [50] to remove Illumina sequencing adapters and poor quality read ends, using the following essential settings: ILLUMINACLIP:2:30:10:2:true, SLIDINGWINDOW:4:10, LEADING:3, TRAILING:3, MINLEN:50. Reads aligning to mitochondrial DNA, ribosomal RNA or phiX174 genome, or composed of a single nucleotide, were removed using STAR (version 2.7.9a) [51]. The remaining reads were aligned to the Gencode human transcriptome version 38 (for genome version hg38) using STAR (version 2.7.9a) [51] with essential non-default settings: --seedSearchStartLmax 12, -- alignSJoverhangMin 15, --outFilterMultimapNmax 100, --outFilterMismatchNmax 33, -- outFilterMatchNminOverLread 0, --outFilterScoreMinOverLread 0.3, and --outFilterType BySJout. The unstranded, uniquely mapping, gene-wise counts for primary alignments were collected in R (version 4.1.0) using Rsubread:feature Counts (version 2.8.1) [52], totaling in 17.0 to 22.6 million per sample. After normalization, using varianceStabilizingTransformation (from DESeq2 version 1.34.0), the data were subjected to sample-level quality control: no obvious batch effects were identified. Differentially expressed genes (DEGs) between experimental groups were identified in R (version 4.2.0) using DESeq2 (version 1.36.0) [53] by employing Wald statistic and lfcShrink for FC shrinkage (type=“apeglm”, version 1.18.0) [54].

### 2.10 Statistical analyses and presentation of data

Data are shown, depending on their distribution, as mean ± SEM or median ± interquartile range as indicated in the figure legends. Statistical analyses were performed using GraphPad Prism9 (version 9.0.0). Normal distribution was tested with the Shapiro-Wilk test. One-way ANOVA (normal distribution) or Kruskal-Wallis test (non-normal distribution) was performed for data with more than two groups and no other variables (*i.e.,* no treatment with Torin 1, Lactacystin, Bafilomycin A1, or Tunicamycin). If a significant difference was observed after the initial ANOVA, this was followed by Tukey’s multiple comparison test (for normally distributed data) or Dunńs multiple comparison test (for not normally distributed data). Two-way ANOVA was performed for data with more than two groups and an additional variable (*i.e.,* treatment with treatment with Torin 1, Lactacystin, Bafilomycin A1, or Tunicamycin). If a significant difference was observed after the initial ANOVA, this was followed by Tukey’s multiple comparison test. p values ≤ 0.05 were considered statistically significant and only significant p values (assessed with the post hoc tests) are indicated in the graphs.

Graphs were drawn using the GraphPad Prism software (version 9.0.0). For Western blot, three independent experiments with cells plated at different passages were considered biological replicates. In the Seahorse assays, results from plating of cells from different passages were considered biological replicates. For quantification of immunofluorescence data (p62), individual pictures, each containing several cells, taken from different areas in the same coverslip were considered biological replicates. The number of n indicated in the figure legends describes the number of biological replicates according to these definitions.

## 3 Results

### 3.1 Fibroblasts from intermediate C9-HRE-carrying iNPH patients show unchanged C9orf72 expression and do not display RNA foci

In this study, we have utilized a cohort of skin fibroblasts obtained from three healthy controlindividuals and four intermediate C9-HRE-carrying and one long C9-HRE-carrying iNPH patient, who later developed ALS during follow-up. It has been suggested in previous studies that the C9-HRE leads to decreased C9orf72 mRNA and protein levels due to haploinsufficiency [28,55]. We therefore first assessed C9orf72 mRNA and protein expression in global RNA sequencing data and protein samples from the fibroblasts of controls and iNPH patients carrying different lengths of the C9-HRE. The mRNA levels were similar in the intermediate or long C9-HRE-carrying iNPH patients to those in controls (Fig. 1 a). Moreover, no significant differences in C9orf72 protein levels were observed between the controls and intermediate or long C9-HRE carriers, even though there was a trend towards increased levels in the intermediate carriers (p = 0.1663) (Fig. 1 b and c). These findings suggest that the intermediate or long C9-HRE-carrying iNPH patient-derived fibroblasts do not display signs of *C9orf72* haploinsufficiency on the mRNA or protein level.

**Fig. 1.**
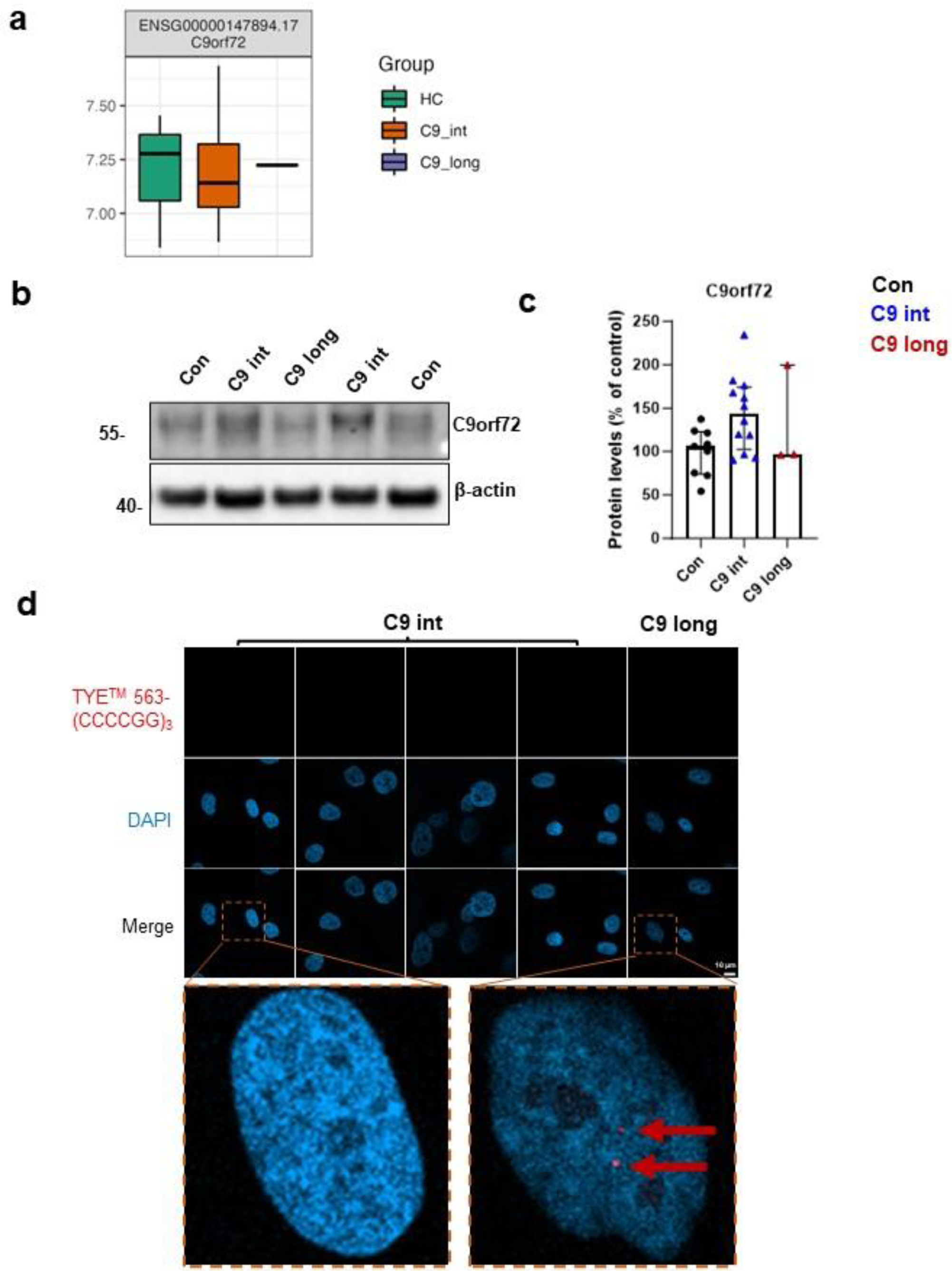
iNPH patient fibroblasts carrying intermediate C9-HRE do not display the main C9-HRE-associated pathological hallmarks. **a)** Quantification of C9orf72 mRNA levels from RNA sequencing data from the fibroblasts of healthy controls (Con) and intermediate (C9 int) and long (C9 long) C9-HRE carriers shows similar levels in all groups. **b)** Representative Western blot images of fibroblast cell lysates from a control (Con), iNPH patient with intermediate C9-HRE (C9 int), and iNPH patient with long C9-HRE (C9 long). The blots were probed with antibodies against C9orf72 and β-actin (loading control used for normalization). **c)** Quantification of the C9orf72 protein levels from the Western blot images. Data are shown as the mean of three separate experiments (=independent platings of cells in different passages) ± SEM. Two-way ANOVA, followed by Tukeýs multiple comparison test, was performed. **d)** Representative images of FISH analysis, revealing RNA foci (red) in the fibroblasts of the C9-HRE long carrier. The intermediate C9-HRE carriers do not show any RNA foci. DAPI (blue) was used to stain the nuclei.

The presence of RNA foci in fibroblasts, cortex, spinal cord, and iPSC-derived skeletal myocytes has been previously described in C9-HRE carriers with ALS and FTD [5,40,41,56].In our previous study, the C9-HRE-carrying FTD patient-derived skin fibroblasts were found to express RNA foci but none of the DPR proteins were detected [42]. Here, FISH analysis indicated that fibroblasts from intermediate C9-HRE-carrying iNPH patients did not display RNA foci but those from the long C9-HRE carrier iNPH patient did (Fig. 1 d), which is in line with previous results [42,56–58].

### 3.2 Fibroblasts of the long but not intermediate C9-HRE-carrying iNPH patient display increased number of p62-positive puncta and do not show alterations in basal or induced autophagy

Since accumulation of p62 has been previously described in the brain of iNPH patients [59] and FTD patients carrying the C9-HRE [30], we examined whether the fibroblasts from iNPH patients showed aggregation of p62. No cytoplasmic inclusions of p62 in the intermediate or long C9-HRE-carrying fibroblasts could be observed (Fig. 2 a), but quantitative analysis of the number of p62-positive puncta revealed a significant increase in the long C9-HRE- carrying iNPH patient fibroblasts compared both to the healthy controls and the intermediate C9-HRE carriers. We also observed a trend towards increased number of puncta in the fibroblasts from the intermediate C9-HRE-carrying iNPH patients compared to healthy controls, but this was not statistically significant (p=0.1563) (Fig. 2 b). Area (Fig. 2 c) or intensity (Fig. 2 d) of the p62-positive puncta did not differ between any of the groups. RNA sequencing data showed that the p62 mRNA levels were not significantly changed between the three groups, indicating that p62 transcription was unaltered (Fig. 2 e). Staining of brain biopsy samples from the same iNPH patients indicated the presence of p62 intracellular inclusions in the long C9-HRE carrier and in the intermediate C9-HRE carrier with the highest number of repeats (23 repeats) (Fig. 2 f)

**Fig. 2.**
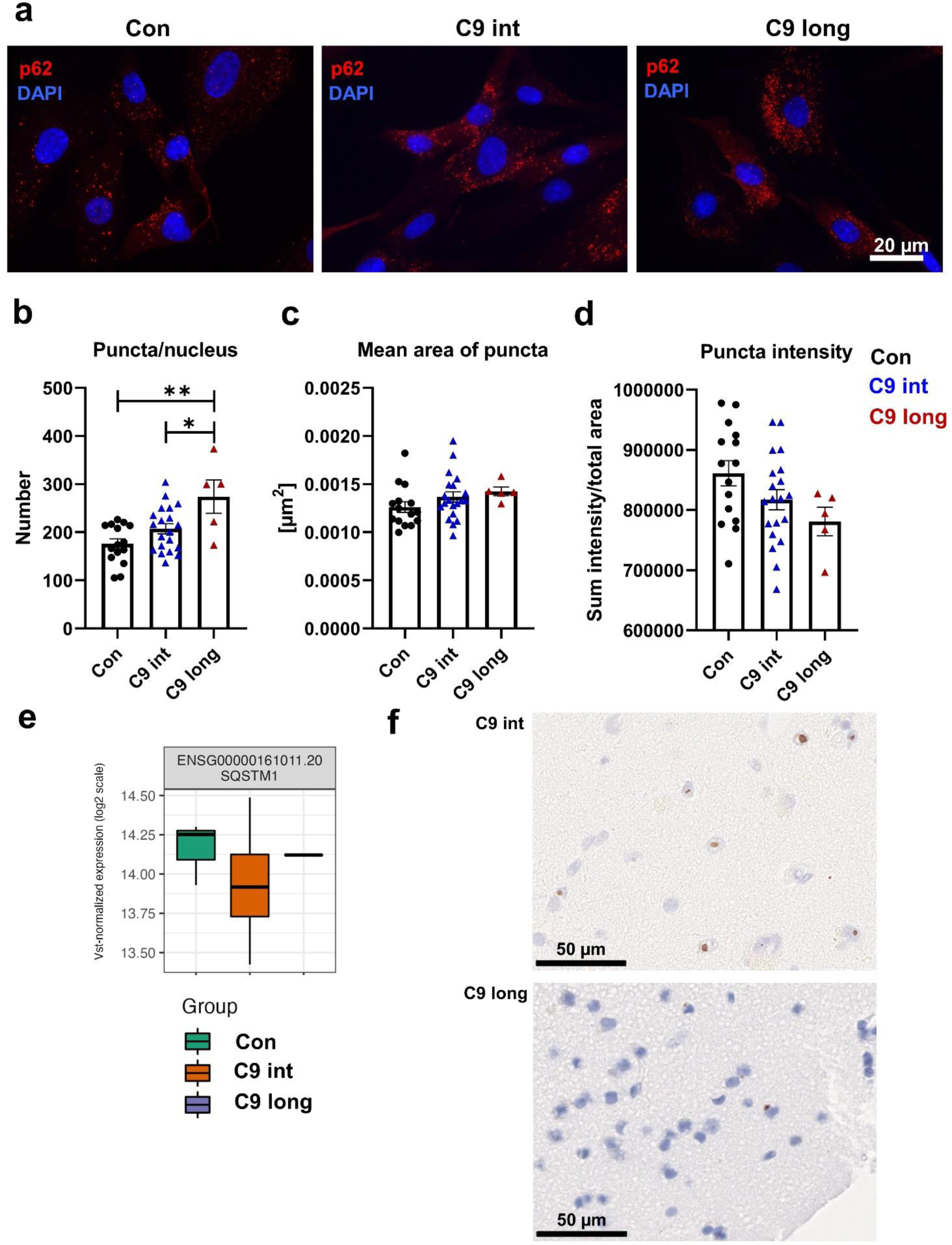
Number of p62-positive puncta is increased in fibroblasts from long but not intermediate C9-HRE carriers. **a)** Representative fluorescence microscopy images of staining with anti-p62 antibody (red) in fibroblasts of a control individual, an iNPH patient with intermediate C9-HRE, and an iNPH patient with long C9-HRE. Nuclei were stained with DAPI (blue). **b)** Quantification of number of p62 puncta. **c)** Quantification of mean area of p62 puncta. **d)** Quantification of intensity of p62 puncta. Data are shown as mean ± SEM and one-way ANOVA, followed by Tukey’s multiple comparison test, was performed for all data sets. Only p values that were significant in the *post hoc* test are indicated in the graphs. Number of images analyzed: n = 15 for control; n = 20 iNPH with intermediate C9-HRE; and n = 5 iNPH with long C9-HRE. Data were obtained from one experiment and each datapoint represents one image. **p ≤ 0.01, ***p ≤ 0.001. **e)** Quantification of mRNA levels of p62/SQSTM1 from RNA sequencing data from healthy controls (Con) and intermediate (C9 int) and long (C9 long) C9-HRE carriers indicates no significant changes between the groups. **f)** Representative microscopy images of p62-positive inclusions (brown) in patient brain biopsies of one intermediate C9-HRE carrier (upper image) and the long C9-HRE carrier (lower image). Nuclei were stained with Haematoxylin (blue).

As p62 is one of the key receptors targeting cargo to the autophagosomes and also itself a substrate for autophagosomal degradation, the increased number of p62-positive puncta could suggest alterations in autophagosomal activity in the iNPH fibroblasts. Moreover, it has been shown that in cortical brain biopsies from iNPH patients, non-fused autophagic vacuoles are more numerous in neuronal somas than in healthy individuals, further implying potentially defective autophagic function [60]. Impaired autophagic function has also been suggested to contribute to the pathogenesis of several other neurodegenerative diseases, including ALS [61]. When autophagy is induced, phosphatidylethanolamine is conjugated to cytosolic LC3BI, which subsequently forms autophagosomal membrane-bound LC3BII. Thus, an increased LC3BII/LC3BI ratio can be used as an indicator of autophagy induction [62].

To first assess basal autophagy in the iNPH fibroblasts, the fibroblasts were treated with BafA1, blocking the late stages of the autophagosomal degradation by inhibiting the fusion of autophagosomes with lysosomes [62–64]. Analysis of the protein levels of LC3BI, LC3BII, and p62 using Western blot, showed a significant increase in the LCBII levels (Fig. 3 a and f) and LC3BII/LC3BI ratio (Fig. 3 d) after the treatment in all fibroblasts. However, no differences in this increase were observed between control and iNPH patient fibroblasts with intermediate or long C9-HRE, suggesting normal basal autophagy in all the iNPH patient-derived fibroblasts. LCBI or p62 levels were similar in all fibroblasts under basal conditions (DMSO) and they remained unchanged with BafA1 treatment (Fig. 3 a, c, and e).

**Fig. 3.**
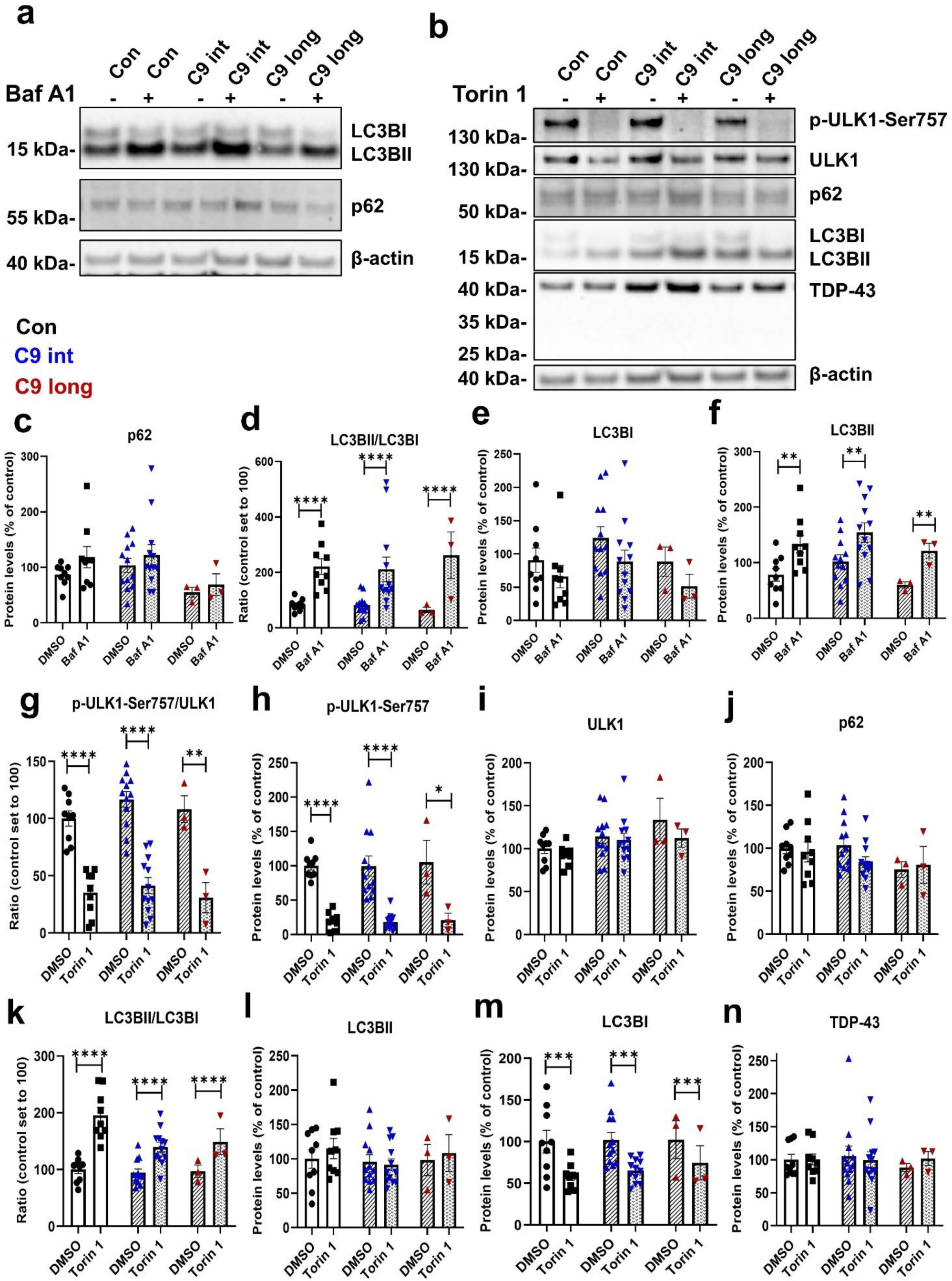
iNPH patient-derived fibroblasts display unaltered basal autophagy and response to an autophagy-inducing stimulus. **a)** Representative Western blot images from fibroblast cell lysates probed with LC3B, p62 and β-actin (loading control for normalization) antibodies. To block the autophagosomal flux and fusion of autophagosomes with lysosomes, fibroblasts were treated with 300 nM bafilomycin (BafA1) for 6 h. DMSO was used as a vehicle control. **b)** Representative Western blot images of p-ULK1-Ser757, ULK1, p62, LC3BI and II, TDP-43, and β-actin from fibroblast cell lysates. Cells were treated with 200 nM Torin 1 overnight to induce autophagy. DMSO was used as a vehicle control. **c-f)** Treatment with BafA1 [**c)** Quantification of p62. **d)** Ratio of LC3BII/I. **e)** Quantification of LC3BI. **f)** Quantification of LC3BII.] **g-n)** Treatment with Torin 1 [**g)** Ratio of p-ULK1-Ser757/ULK1. **h)** Quantification of p-ULK1-Ser757. **i)** Quantification of ULK1. **j)** Quantification of p62. **k)** Ratio of LC3BII/I. **l)** Quantification of LC3BII. **m)** Quantification of LC3BI. **n)** Quantification of TDP-43]. Data are shown as the mean of three separate experiments (=independent platings of cells in different passages) ± SEM. Two-way ANOVA, followed by Tukey’s multiple comparison test, was performed for all data sets. Only p values that were significant in the *post hoc* test are indicated in the graph. n = 9 control; n = 12 iNPH with intermediate C9-HRE; and n = 3 iNPH with long C9-HRE. *p ≤ 0.05, **p *≤* 0.01, ***p ≤ 0.001, ****p ≤ 0.0001.

Pharmacological induction of autophagy may uncover defects in autophagy even when the basal autophagy is not impaired [65]. To examine this, we induced autophagy with Torin 1 and assessed the protein levels of the autophagy-associated proteins ULK1, phospho-ULK1 (p-ULK1-Ser757), LC3BI and II, and p62, and also TDP-43 (Fig. 3 b and g-n). ULK1 levels were similar in all the fibroblasts in basal conditions (DMSO) and after Torin 1 treatment (Fig. 3 b and i). As expected, treatment with Torin 1 significantly decreased the p-ULK1-Ser757 levels (Fig. 3 b and h) and the ratio of p-ULK1-Ser757 to ULK1, indicating induction of autophagy. However, no differences could be observed between the fibroblasts from iNPH patients carrying the intermediate or long C9-HRE and healthy controls (Fig. 3 b and g). Treatment with Torin 1 did not affect the LC3BII levels (Fig. 3 b and l), but significantly decreased the LC3BI levels (Fig. 3 b and m), leading to a significantly increased LCBII to LC3BI ratio. Again, no differences were observed between the iNPH patient and control fibroblasts (Fig 3. b and k), suggesting that iNPH patient-derived fibroblasts can respond normally to an autophagy-inducing stimulus. TDP-43 levels were similar in all the fibroblasts and remained unaltered after Torin 1 treatment (Fig. 3 b and n). No TDP-43 C-terminal fragments were detected in any of the fibroblasts (Fig. 3 b). Taken together, modulating different phases of the autophagosomal degradation pathway indicate that the iNPH fibroblasts do not show deficits in autophagy.

### 3.3 Fibroblasts from intermediate or long C9-HRE-carrying iNPH patients display unaltered proteasomal activity and subcellular localization and phosphorylation of TDP-43

Not only impaired autophagy but also defects in UPS have been suggested to contribute to pathological protein aggregation in neurodegenerative diseases. To assess UPS function in iNPH fibroblasts, we blocked the UPS with the proteasomal inhibitor lactacystin and examined the levels of poly-ubiquitinated proteins, as well as TDP-43 and p-TDP-43, which typically show pathological accumulation in FTD brain [40]. As expected, lactacystin treatment significantly increased the level of poly-ubiquitinated proteins, but there were no differences between healthy control and iNPH patient-derived fibroblasts (Fig. 4 b and c). The levels of p-TDP-43 (Fig. 4 b and d) and TDP-43 (Fig. 4 b and e) were also similar in healthy controls and iNPH patient-derived fibroblasts and treatment with lactacystin did not affect their levels. Thus, also the p-TDP-43/TDP-43 ratio remained unaltered (Fig. 4 f). According to RNA sequencing, there were no significant differences in the TARDBP mRNA levels between the three groups (Fig. 4 a).

**Fig. 4.**
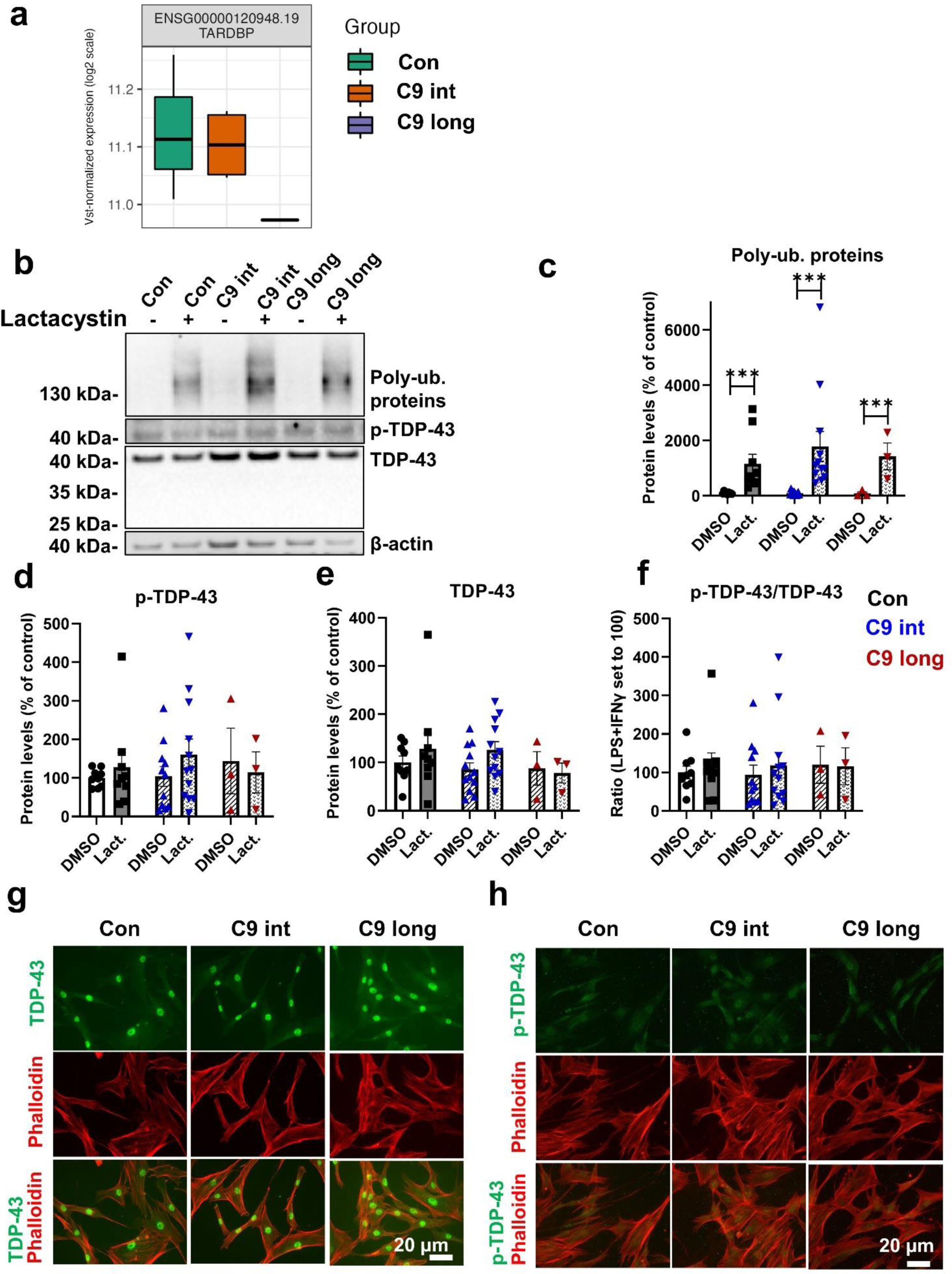
Levels or subcellular localization of TDP-43 and p-TDP-43 are not affected in iNPH patient-derived fibroblasts. **a)** Quantification of TARDBP mRNA levels from RNA sequencing data from healthy controls (Con) and intermediate (C9 int) and long (C9 long) C9-HRE carriers shows similar levels in intermediate C9 HRE carriers and controls. **b)** Representative Western blot images from fibroblast cell lysates probed with antibodies against poly-ubiquitinated (Poly-Ub.) proteins, p-TDP-43, TDP-43, and β-actin (loading control for normalization) antibodies. Cells were treated with 10 µM lactacystin for 16 h to block protein degradation through the UPS. DMSO was used as a vehicle control. **c)** Quantification of Poly-Ub. proteins. **d)** Quantification of p-TDP-43. **e)** Quantification of TDP-43. **f)** Ratio of p-TDP-43/TDP-43. Data are shown as the mean of three separate experiments (=independent platings of cells in different passages) ± SEM. Two-way ANOVA, followed by Tukey’s multiple comparison test, was performed. Only p values that were significant in the *post hoc* test are indicated in the graphs. n = 9 control; n = 12 iNPH with intermediate C9-HRE; and n = 3 iNPH with long C9-HRE. ***p ≤ 0.001. **g)** Representative fluorescence microscopy images of staining with anti-TDP-43 antibody (green) and Phalloidin (red) in fibroblasts of control (left column), iNPH patient with intermediate C9-HRE (middle column), and iNPH patient with long C9-HRE (right column). Neither changes in subcellular localization nor formation of TDP-43-positive inclusions could be observed. **h)** Representative fluorescence microscopy images of staining with anti-p-TDP-43 antibody (green) and Phalloidin (red) in fibroblasts of control (left column), iNPH patient with intermediate C9-HRE (middle column), and iNPH patient with long C9-HRE (right column). Neither changes in subcellular localization nor formation of p-TDP-43 positive inclusions could be observed. Images were taken from one experiment.

The RNA-binding protein TDP-43 can shuttle between the nucleus and cytoplasm [66] and cytoplasmic accumulation has been observed in the CNS of ALS and FTD patients carrying the C9-HRE [31]. We therefore wanted to assess whether changes in subcellular localization of TDP-43 and p-TDP-43 could be observed. TDP-43 was strongly localized in the nucleus in all fibroblasts with no discernible differences between the groups (Fig. 4 g). p-TDP-43 showed both nuclear and cytoplasmic subcellular localization (Fig. 4 h) but there were no apparent differences between iNPH patient-derived fibroblasts and controls. No cytoplasmic inclusion bodies containing TDP-43 or p-TDP-43 were observed in any of the fibroblasts (Fig. 4 g and h).

### 3.4 iNPH patient-derived fibroblasts carrying intermediate and long C9-HRE show altered energy metabolism

Mitochondrial respiration and glycolysis are the main energy-producing pathways in cells and impaired energy metabolism has been described in several neurodegenerative diseases, including AD, PD, ALS, and FTD [67,68]. To assess mitochondrial function, we examined energy metabolism of the fibroblasts related to oxidative phosphorylation by measuring changes in OCR after treatment with different ETC modulators in control and iNPH fibroblasts (Fig. 5 a). A significant reduction in the basal respiration in iNPH fibroblasts carrying the long C9-HRE compared to controls could be observed (Fig. 5 b) with a similar, but non-significant, trend (p=0.079) in the intermediate C9-HRE carrier iNPH patient fibroblasts. Moreover, respiration linked to ATP production (Fig. 5 e) was significantly reduced in the fibroblasts of the intermediate C9-HRE carriers and an even stronger reduction could be observed in the fibroblasts with the long C9-HRE, suggesting impaired mitochondrial function. Also, non-mitochondrial respiration (Fig. 5 g) was significantly reduced in the fibroblasts with the long C9-HRE, which, together with the observed deficits in the other components of the mitochondrial respiratory chain, might indicate an overall decrease in the energy metabolism of fibroblasts with the long C9-HRE. Maximal respiration (Fig. 5 c), spare capacity (Fig. 5 d), and proton leak (Fig. 5 f) were similar in iNPH patient-derived fibroblasts and controls.

**Fig. 5.**
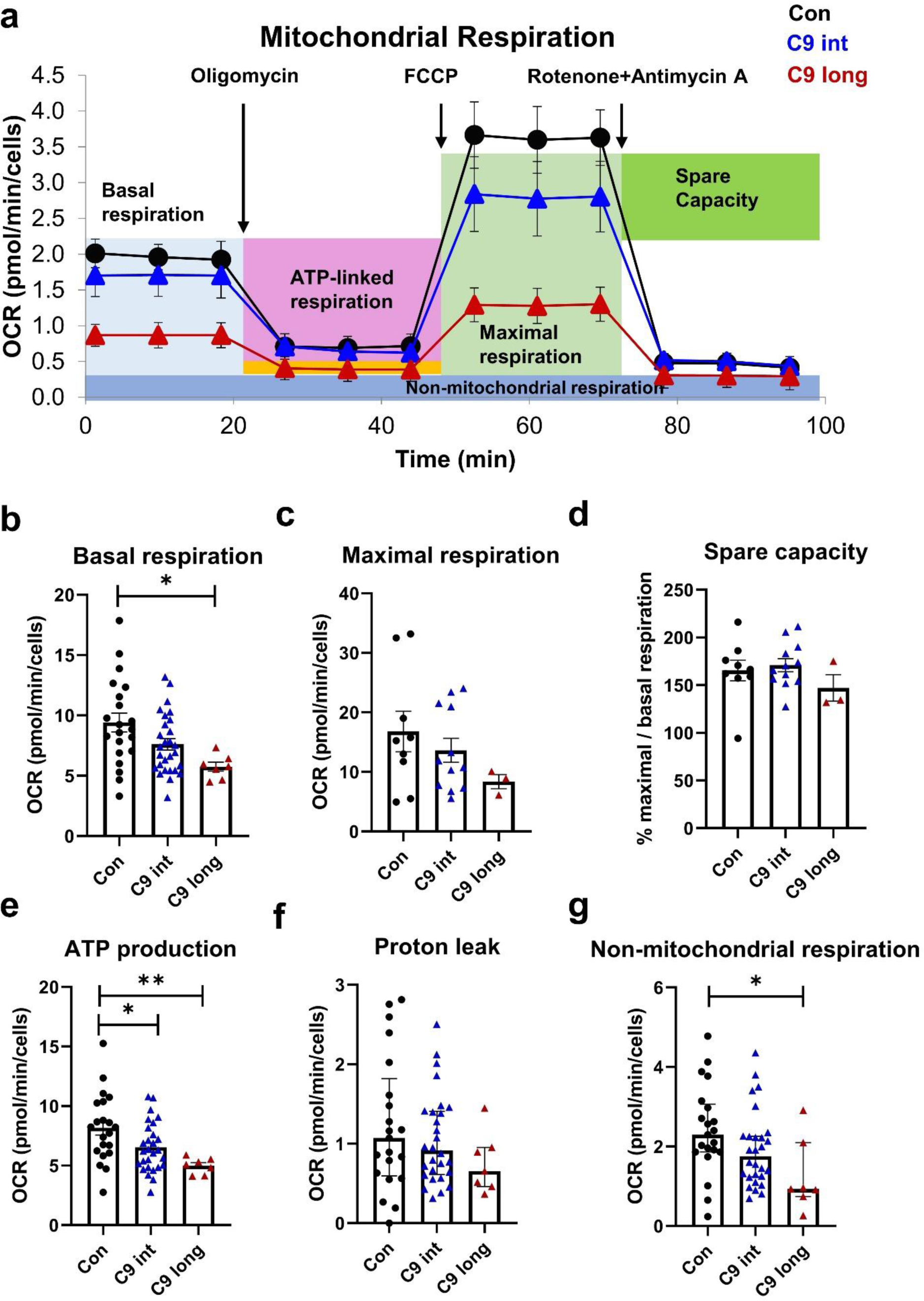
Mitochondrial respiration is impaired in iNPH patient-derived fibroblasts from both intermediate and long C9-HRE carriers. Using the Cell Mito Stress Test, several parameters of mitochondrial function were assessed. **a)** Example of Cell Mito Stress Test in fibroblasts of one control individual, one iNPH patient with intermediate C9-HRE, and one iNPH patients with long C9-HRE fibroblast line. **b)** Quantification of basal respiration. **c)** Quantification of maximal respiration. **d)** Quantification of spare capacity. **e)** Quantification of ATP production. **f)** Quantification of proton leak. **g)** Quantification of non-mitochondrial respiration. Data are shown as the mean ± SEM, and one-way ANOVA, followed by Tukey’s multiple comparison test, was performed (b-e). Data are shown as median ± interquartile range and Kruskal-Wallis, followed by Dunńs multiple comparison test, was performed (f, g). Only p values that were significant in the *post hoc* test are indicated in the graphs. n = 9 control; n = 12 iNPH with intermediate C9-HRE; and n = 3 iNPH with long C9-HRE for maximal respiration and spare capacity and n = 21 control; n = 28 iNPH with intermediate C9-HRE; and n = 7 iNPH with long C9-HRE for other parameters. *p *≤* 0.05; **p *≤* 0.01. Abbreviations: FCCP, cyanide-4-(trifluoromethoxy)phenylhydrazone; OCR, oxygen consumption rate.

To assess glycolytic function, we investigated changes in ECAR after treatment with glucose and oligomycin in control, intermediate, and long C9-HRE iNPH fibroblasts (Fig. 6 a). Interestingly, a significant increase in glycolysis could be observed in the fibroblasts of the intermediate C9-HRE-carrying iNPH patients as compared to control fibroblasts. The difference compared to the long C9-HRE iNPH fibroblasts did not reach statistical significance (p=0.994) and there was no significant difference in glycolytic activity between the long C9-HRE carrier and the controls (Fig. 6 b). However, the glycolytic capacity of the iNPH fibroblasts from the long C9-HRE carrier was significantly decreased when compared to the intermediate C9-HRE carriers (Fig. 6 c). There was also a trend towards a decreased glycolytic reserve in the long C9-HRE-carrying iNPH patient fibroblasts when compared to control fibroblasts, but this decrease did not reach statistical significance (p=0.0633). Interestingly, similar results were obtained in the fibroblasts from FTD patients carrying the long C9-HRE. In these cells, glycolysis and glycolytic capacity were significantly reduced compared to intermediate C9-HRE carriers with iNPH (Supplementary Fig. 1).

**Fig. 6.**
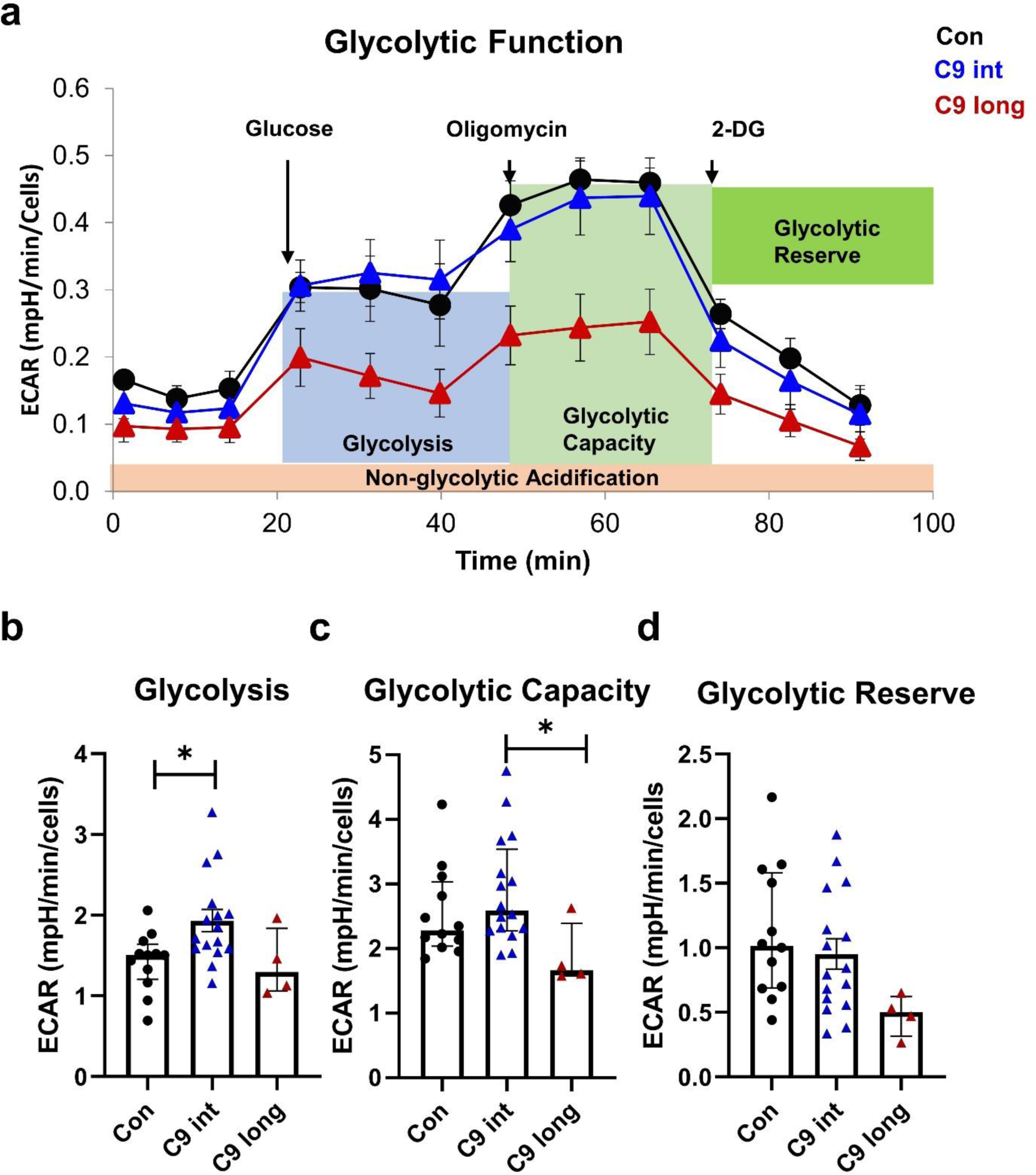
Glycolysis is differently affected in intermediate and long C9-HRE-carrying iNPH patient-derived fibroblasts. Using the Cell Glycolysis Stress Test, several parameters of glycolysis were assessed. a) Example of Cell Glycolysis Stress Test in fibroblasts of one control individual, one intermediate C9-HRE carrier iNPH patient, and one long C9-HRE iNPH patient fibroblast line. b) Quantification of glycolysis. c) Quantification of glycolytic capacity. d) Quantification of glycolytic reserve. Data are shown as mean ± SEM, and one-way ANOVA, followed by Tukey’s multiple comparison test, was performed (b, d). Data are shown as median ± interquartile range and Kruskal-Wallis, followed by Dunńs multiple comparison test, was performed (c). Only p values that were significant in the *post hoc* test are indicated in the graphs. n = 12 control; n = 16 iNPH with intermediate C9-HRE; and n = 4 iNPH with long C9-HRE from four separate experiments. *p ≤ 0.05; **p ≤ 0.01. Abbreviations: 2-DG; 2-deoxy-D-glucose ECAR, extracellular acidification rate

### 3.5 Intermediate C9-HRE-carrying fibroblasts show only moderate gene expression changes as compared to healthy control fibroblasts

To assess potential gene expression changes in the intermediate C9-HRE-carrying fibroblasts, global RNA sequencing analysis was performed. Only 32 DEGs were identified and of these 16 were downregulated and 16 upregulated in the intermediate C9-HRE-carrying fibroblasts as compared to control fibroblasts (Fig. 7 and Supplementary Fig. 2). This analysis did not reveal specific gene expression changes related to autophagy, UPS, or mitochondrial energy metabolism, except for the upregulation of *CTP1C*. This gene encodes carnitine palmitoyltransferase 1C protein, which regulates the beta-oxidation and transport of long-chain fatty acids into mitochondria, and thus may play a role in the regulation of energy homeostasis related to ATP and NADPH production. Such function for *CTP1C* has been described *e.g.* in cancer [69]. One previous report on brain transcriptome data has shown decreased expression of *CPT1C* in the cerebellum but not the frontal cortex of patients with the C9-HRE [70].

**Fig. 7.**
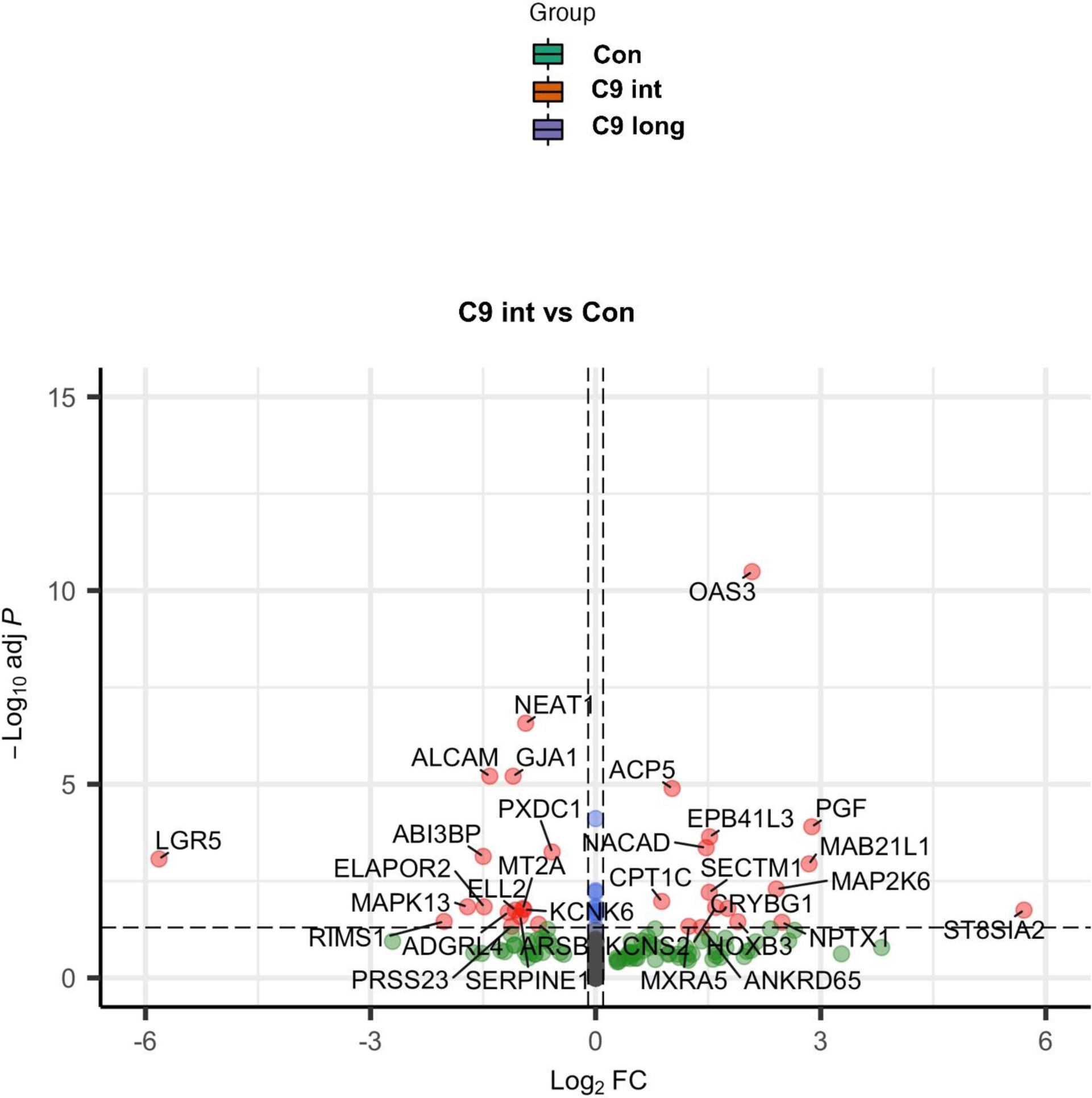
Differentially expressed genes in intermediate C9-HRE-carrying iNPH patient-derived fibroblasts compared to healthy control fibroblasts. Volcano plot showing the differentially expressed genes (DEGs). Altogether 16 genes were significantly upregulated and 16 significantly downregulated in the fibroblasts from the intermediate C9-HRE-carrying iNPH patients as compared to those from healthy controls.

## 4 Discussion

In this study, our aim was to assess whether carrying the intermediate C9-HRE leads to the development of cellular pathologies similar to those with long C9-HRE and whether it affects cellular functions related to protein degradation and energy metabolism in skin biopsy-derived fibroblasts of iNPH patients.

Haploinsufficiency is one of the mechanisms associated with the C9-HRE, indicated by decreased levels of C9orf72 mRNA and protein expression, which have been observed in patient CNS and also in the periphery, for example in lymphocytes and blood [5,7,55]. Here, we did not find significant changes in C9orf72 mRNA or protein levels in the intermediate carriers or the long C9-HRE carrier, which is in line with our previous results in fibroblasts from long C9-HRE-carrying FTD patients [42], suggesting that fibroblasts with intermediate or long C9-HRE do not display haploinsufficiency. Increased C9orf72 expression has been observed in the brain tissue of intermediate C9-HRE carriers [25], but in the present study, only a trend towards increased C9orf72 protein levels could be observed in the intermediate C9-HRE fibroblasts.

Nuclear RNA foci, a C9-HRE-associated gain-of-toxic-function hallmark, were observed in the fibroblasts of the long C9-HRE carrier but not the intermediate carriers. This is in agreement with previous publications on fibroblasts from C9-HRE carriers suggesting that carriers of intermediate repeats shorter than 30 do not develop this pathology [42,56]. Data from patients with corticobasal degeneration carrying intermediate C9-HRE further support this finding as no RNA foci could be found in the brain or spinal cord of these patients [25].

A trend towards an increased number of p62-positive puncta in the fibroblasts of the intermediate C9-HRE carriers was observed in the present study, but this was not statistically significant. In contrast, fibroblasts of the long C9-HRE carrier, showed a significant increase in the number of p62-containing puncta, in line with our previous study in the fibroblasts of FTD patients carrying the long C9-HRE, which showed a significant increase in number, size, and intensity of p62-containing vesicles [42]. p62-containing inclusion bodies were not detected in this or our previous study. p62 pathology is observed in the brain of FTD spectrum patients and the DPR proteins have been shown to co-localize in p62-positive inclusions in the brains of C9-HRE carriers [26]. Interestingly, p62 brain pathology has been previously observed in an iNPH patient in a case report [59] and also in brain biopsies from the patient with the long C9-HRE and in one of the intermediate C9-HRE carriers in this study. These results indicate that similarly to the brain, although not forming intracellular inclusions, skin fibroblasts, especially from the long C9-HRE carriers, show accumulation of the p62 protein. This could suggest impaired protein degradation via the autophagosomal or proteasomal pathway, as p62 can undergo degradation through either of these pathways.

However, in detailed analyses of the protein degradation pathways, we did not observe changes in the basal or induced autophagy in the fibroblasts of the intermediate or long C9-HRE carriers, which is in line with our previous work on FTD patient-derived fibroblasts, where no changes in the autophagosomal pathway were observed [42]. The finding of unaltered autophagy also agrees with the unchanged C9orf72 protein levels in the C9-HRE-carrying fibroblasts. Regulation of autophagy by the C9orf72 protein isoform A has been suggested in several studies but it is still controversial whether the reduction of the C9orf72 protein levels leads to increased or decreased autophagy in different cell types [28,43,48,63,65,71–75]. The unaltered UPS function in the C9-HRE intermediate or long carriers is also in accordance with our previously published results on FTD patient-derived fibroblasts [42,76,77]. These results indicating normal function of the autophagosomal and proteasomal pathways further suggest that impaired degradation of the p62 protein does not likely cause the observed accumulation of p62 vesicles in the C9-HRE-carrying fibroblasts. Increased levels of p62, without significant changes in LC3 II/LC3-I turnover, have been observed in iPSC-motor neurons of ALS and FTD patients [78], pointing towards other possible underlying mechanisms than autophagy. One alternative mechanism could be increased transcription of the p62 mRNA [79–81]. Our RNA sequencing data, however, suggested that the p62 mRNA levels do not significantly differ between the healthy controls and iNPH patient-derived fibroblasts carrying either intermediate or long C9-HRE, rendering the mechanism underlying the accumulation of the p62-positive vesicles elusive. Moreover, we did not observe TDP-43 or p-TDP-43 mislocalization or aggregation, and their levels were similar and not affected by proteasomal inhibition in healthy control and iNPH patient-derived fibroblasts carrying either intermediate or long C9-HRE. Some previous studies have reported hyperphosphorylation and altered subcellular localization of TDP-43 in patient fibroblasts carrying C9-HRE or other mutations or after proteasomal inhibition using MG132 [82–84]. These studies suggest that significant changes in TDP-43 and p-TDP-43 localization, levels, or aggregation might be dependent on the prevailing stress condition.

Pathological alterations in mitochondria have been observed in neurons from cortical brain biopsies of iNPH patients, indicated by altered numbers of mitochondria-endoplasmic reticulum contact sites [60] and changes in mitochondrial morphology [85]. Mitochondrial dysfunction in iNPH-patient derived fibroblasts has not been studied before but has been observed in the fibroblasts of FTD and ALS patients carrying the C9-HRE [42,47]. Here, we detected altered mitochondrial function in fibroblasts from both intermediate and long C9-HRE-carriers compared to healthy controls. Basal respiration was significantly reduced in the long C9-HRE carrier with a similar trend in the intermediate C9-HRE carriers. Respiration linked to ATP production was significantly reduced in both intermediate and long C9-HRE carriers, similarly to our previous study in long C9-HRE-carrying FTD patient-derived fibroblasts [42]. The detected decrease in non-mitochondrial respiration in the long C9-HRE carrier iNPH fibroblasts could indicate an overall reduction in the energy metabolism of these cells. It is interesting to note that the intermediate C9-HRE carriers showed some impairments in mitochondrial function, although these appeared milder than those in the long C9-HRE carriers.

In the intermediate C9-HRE carriers, the global RNA sequencing showed upregulation of one gene related to mitochondrial function, the *CTP1C* gene, which encodes for the carnitine palmitoyltransferase 1C protein (CPT-1C). A study in iPSC-derived microglia [86] has shown that TREM2 activation and subsequent increase in different acyl carnitine species increases mitochondrial function, whereas treatment with the CPT-1 inhibitor etomoxir abolished this effect. This observation suggests that upregulation CPT-1 might improve mitochondrial function. Thus, the increased expression of *CTP1C* observed in our study in the intermediate C9-HRE carriers might be an attempt of the cells to potentially alleviate the impaired mitochondrial function.

Surprisingly, glycolysis, the other major source of ATP for energy production, was significantly increased in the fibroblasts of intermediate C9-HRE carriers when compared to controls. This might represent a potential compensatory mechanism to counteract the impaired mitochondrial respiration. This idea could be supported by the previous findings showing that fibroblasts from ALS patients carrying the superoxide dismutase 1 mutation (*SOD1*) as well as neuronal NSC-34 cells expressing mutant *SOD1* have reduced mitochondrial respiration but upregulated glycolysis to better meet the ATP demand [45,87]. Moreover, in a study using mouse embryonic fibroblasts isolated from *C9orf72* knockout mice, a significant increase in glycolytic activity could be observed [88]. A similar increase in glycolytic activity could not be observed in the long C9-HRE carrier fibroblasts and they, in fact, showed reduced glycolytic capacity, suggesting that they cannot respond to an increased energy demand as well as the fibroblasts from intermediate C9-HRE carriers and healthy controls. This might contribute to a more pronounced deficit in the overall energy metabolism, involving impairments in both mitochondrial respiration and glycolysis in the long C9-HRE-carrying iNPH fibroblasts. In line with these findings, we also observed similarly impaired glycolytic function in the fibroblasts of FTD patients carrying the long C9-HRE, further underlining the more severely impaired energy production in fibroblasts carrying the long C9-HRE. The findings that the intermediate C9-HRE-carrying fibroblasts do not show evident cell pathologies or drastic functional deficits in the protein degradation pathways or energy metabolism are in concordance with the gene expression data, showing only modestly altered gene expression as compared to the healthy control fibroblasts. Thus, based on these results and current data in the literature, it appears that the intermediate C9-HREs are not highly pathogenic, but might predispose cells for the development of cellular pathologies or deficient protein degradation or energy metabolism under stress conditions.

## 5 Conclusions

The results from the present study demonstrate that iNPH patient-derived fibroblasts carrying the intermediate C9-HRE do not express RNA foci. Neither they nor the long C9-HRE show haploinsufficiency. While the fibroblasts carrying the long C9-HRE exhibit accumulation of p62-positive vesicles, in line with p62-positive inclusions detected in a brain biopsy from the same patient, we did not detect any alterations in the proteasomal or autophagosomal pathways. This suggests that other, yet unknown mechanisms could be responsible for p62 accumulation in the fibroblasts. Nevertheless, these findings indicate that the skin fibroblasts may show similar cell pathologies to those in the brain of the long C9-HRE carriers. The energy metabolism, especially the mitochondrial respiration, is impaired in fibroblasts from both the intermediate and long C9-HRE carriers, but this impairment is more severe in the long C9-HRE-carrying fibroblasts. Taken together, our data suggest that in addition to brain cells, skin fibroblasts can be utilized to investigate some of the underlying disease mechanisms and cell pathologies related to the C9-HRE. The skin fibroblasts might also prove useful and more easily accessible and manageable patient-derived cells for future biomarker discovery and drug testing compared to, for instance, the iPSC-based brain cells.

## List of abbreviations

ALS: amyotrophic lateral sclerosis
ATP: Adenosine triphosphate
BafA1: bafilomycin A1
BSA: bovine serum albumin
bvFTD: behavioral variant frontotemporal dementia
C9 HRE: hexanucleotide repeat expansion in the C9orf72 gene
CNS: central nervous system
DEPC: diethyl pyrocarbonate
ETC: electron transport chain
FCCP: carbonyl cyanide-4-(trifluoromethoxy)phenylhydrazone
FISH: Fluorescence in situ hybridization
FTD: Frontotemporal dementia
GRN: Granulin
IMDM: Iscove’s Modified Dulbecco’s Medium
iNPH: idiopathic normal pressure hydrocephalus
iPSC: induced pluripotent stem cells
LC3B: Microtubule-associated protein 1 light chain-3 B
MAPT: microtubule-associated protein tau
OCR: oxygen consumption rate
p62: sequestosome-1/ ubiquitin-binding protein p62
PFA: paraformaldehyde
SOD1: Superoxide dismutase 1
TARDBP: TAR DNA binding protein
TBST: Tris-buffered saline with 0.1% Tween 20
TDP-43: TAR DNA-binding protein-43
UPS: ubiquitin-proteasome system
VCP: valosin-containing protein

## Acknowledgements

We are grateful to Drs. Jari Koistinaho and Šárka Lehtonen (A.I. Virtanen Institute for Molecular Sciences (AIVI), UEF, Kuopio, Finland and HiLIFE, University of Helsinki, Helsinki, Finland) for sharing their kind help with the establishment of the human skin fibroblast cultures. UEF Cell and Tissue Imaging Unit is acknowledged for providing LSM800 training and facilities. We wish to thank Mr. Ulrich Rostalski for kindly providing user-friendly Excel macros that helped to analyze the microscopy data. This study is part of the research activities of the Finnish FTD Research Network (FinFTD).

## Statements and Declarations

### Funding

This study was supported by the Academy of Finland, grant nos. 315459 (AH), 338182 (MH), 315460 (AMP); 339767 (VL); Sigrid Jusélius Foundation (AH, MH, ES, VL); Finnish Brain Foundation (RW); Orion Research Foundation (HR, RW); Finnish Medical Foundation (ES); Kuopio University Hospital VTR Fund (ES, VL); the Strategic Neuroscience Funding of the University of Eastern Finland (AH, MH); The Maud Kuistila Memorial Foundation (HR); Kuopio University Foundation (HR); ALS tutkimuksen tuki ry. registered association (HR, SL); and the Alfred Kordelin Foundation (RW). HR, SL, NH, and RW were supported by the University of Eastern Finland (UEF) Doctoral Programs in Molecular Medicine (DPMM), and GenomMed. This publication is part of a project that has received funding from the European Union’s Horizon 2020 research and innovation program under the Marie Skłodowska-Curie grant agreement no 740264.

### Competing interests

The authors have no conflicts of interest to declare that are relevant to the content of this article.

### Author contribution

DH, NH, HR, TH, SL, TR, and RW performed the experiments. DH analyzed the data and performed statistical analyses. SH performed the bioinformatic analyses of the RNA sequencing data. PH and VL performed the skin biopsies of the participating individuals. VK genotyped the fibroblast samples for the presence or absence of the C9-HRE. DH and AH wrote the first manuscript draft. ES, AMP, VL, and MH contributed to the study design, supervision, and interpretation of the data. DH and AH conceived the study and research design. AH obtained the main funding supporting the study and supervised all aspects of the study. All authors read and approved the final manuscript.

### Availability of data and material (data transparency)

All data generated or analyzed during this study are included in this published article.

### Ethics approval and consent to participate

All the participants gave a written informed consent. The research in human subjects was performed in accordance with the ethical standards of Declaration of Helsinki and approved by the Research Ethics Committee of the Northern Savo Hospital District (currently: Medical Research Ethics Committee of Wellbeing Services County of North Savo), Kuopio, Finland (ethical permits 16/2013 and 254/2015). Studies on FTD patient-derived skin fibroblasts were performed with the permission 123/2016 and iNPH patient-derived skin fibroblasts with 276/2016 from the Research Ethics Committee of the Northern Savo Hospital District.

### Consent for publication

Not applicable.

## Supplementary Figures

**Supplementary Fig. 1.**
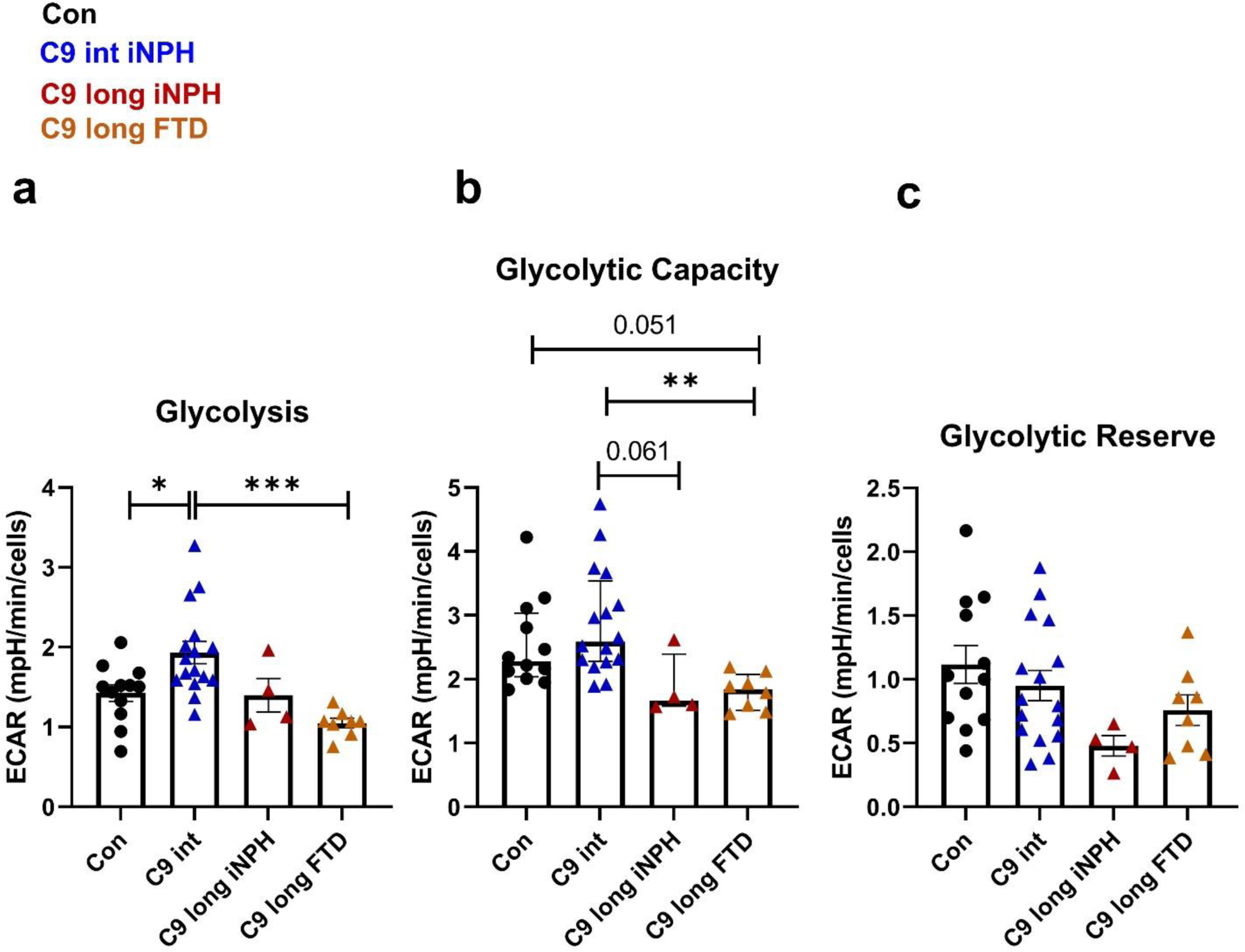
Fibroblasts derived from iNPH and FTD patients show similar impairments in glycolysis. Using the Cell Glycolysis Stress Test, several parameters of glycolysis were assessed. **a)** Glycolysis and **b)** glycolytic capacity in two long C9-HRE carriers with FTD are similarly impaired as in the long C9-HRE carrier with iNPH. **c)** The long C9-HRE-carrying FTD and iPNH patients also show a mild, but non-significant trend towards decreased glycolytic reserve. Data are shown as mean ± SEM, and one-way ANOVA, followed by Tukey’s multiple comparison test, was performed Only p values that were significant in the *post hoc* test are indicated in the graphs. n = 12 control; n = 16 iNPH with intermediate C9-HRE; n = 4 iNPH with long C9-HRE; n = 8 FTD with long C9-HRE. *p ≤ 0.05; **p ≤ 0.01, ***p ≤ 0.001. Abbreviations: ECAR, extracellular acidification rate.

**Supplementary Fig. 2.**
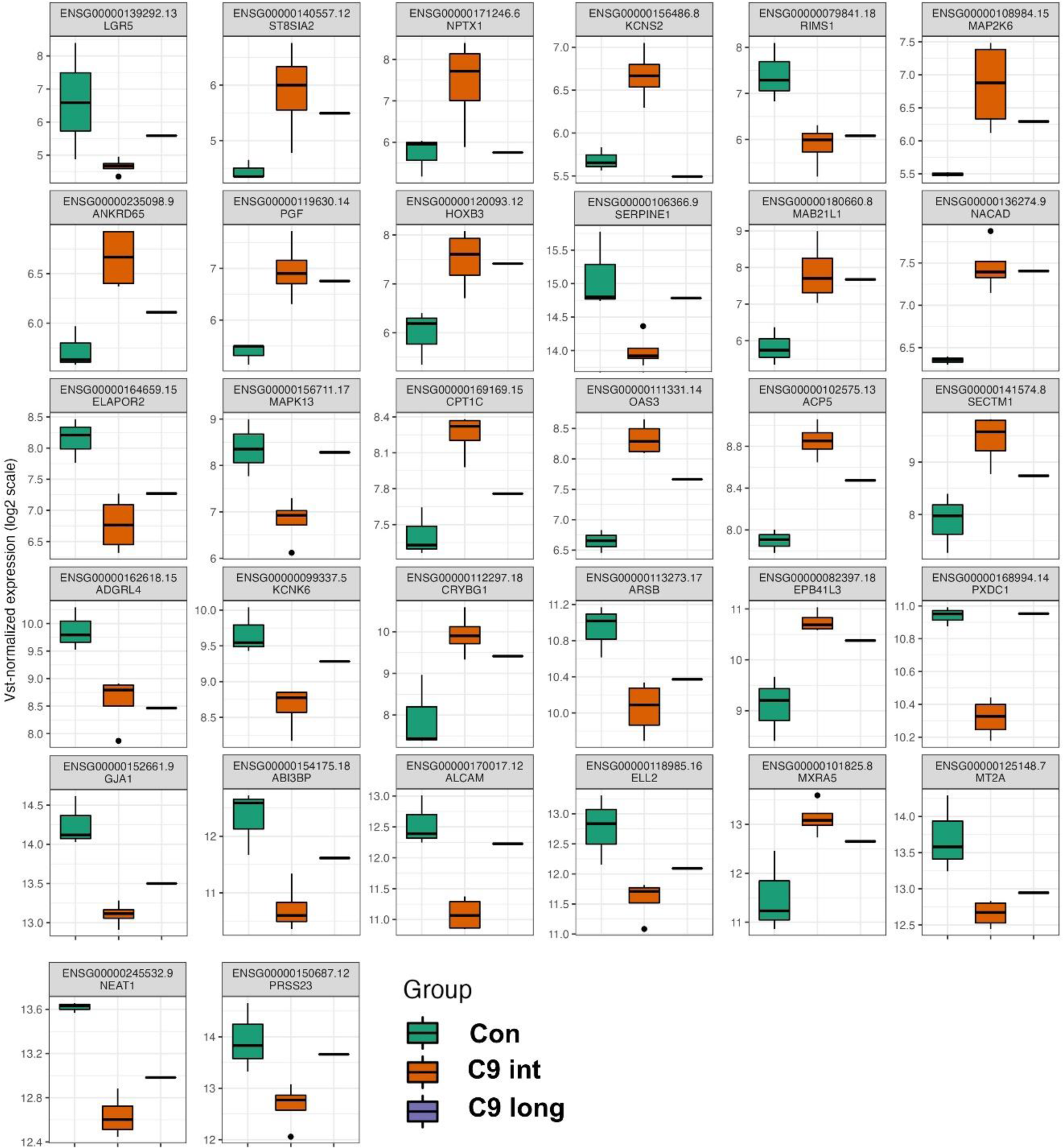
Differentially expressed genes in fibroblasts from healthy controls and intermediate and long C9-HRE-carrying iNPH patients. In total, 32 differentially expressed genes (DEGs) with statistically significant expressional changes were identified. Of these 16 were upregulated and 16 downregulated in the intermediate C9-HRE carrier (C9 int) compared to healthy control (Con) fibroblasts. Data from one long C9-HRE carrier (C9 long) fibroblasts are shown in the boxplots but not included in the statistical analyses.

## Notes

### Competing Interest Statement

The authors have declared no competing interest.

